# Simultaneous Modelling of Movement, Measurement Error, and Observer Dependence in Double-Observer Distance Sampling Surveys

**DOI:** 10.1101/126821

**Authors:** By Paul B. Conn, Ray T. Alisauskas

## Abstract

Mark-recapture distance sampling uses detections, non-detections and recorded distances of animals encountered in transect surveys to estimate abundance. However, commonly available distance sampling estimators require that distances to target animals are made without error and that animals are stationary while sampling is being conducted. In practice these requirements are often violated. In this paper, we describe a marginal likelihood framework for estimating abundance from double-observer data that can accommodate movement and measurement error when observations are made consecutively (as with front and rear observers) and when animals are randomly distributed when detected by the first observer. Our framework requires that two observers independently detect and record binned distances to observed animal groups, as we well as a binary indicator for whether animals were moving or not. We then assume that stationary animals are subject to measurement error whereas moving animals are subject to both movement and measurement error. Integrating over unknown animal locations, we construct a marginal likelihood for detection, movement, and measurement error parameters. Estimates of animal abundance are then obtained using a modified Horvitz-Thompson-like estimator. In addition, unmodelled heterogeneity in detection probability can be accommodated through observer dependence parameters. Using simulation, we show that our approach yields low bias compared to approaches that ignore movement and/or measurement error, including in cases where there is considerable detection heterogeneity. We demonstrate our approach using data from a double-observer waterfowl helicopter survey.

## 1. Introduction

Distance sampling surveys (Burnham, Anderson and Laake, 1980; Buckland et al., 2001) are often used to estimate the abundance of wildlife populations. Historically, such surveys were conducted by a single observer who followed a transect line and recorded the perpendicular distance to each detected animal group. Assuming 100% detection on the transect line, models can be fitted to these data that estimate abundance over the surveyed area while accounting for detection probabilities that decline with distance from the transect line.

More recently, investigators discovered that double-observer surveys have some large advantages over single-observer surveys. For instance, one can use records of detection/non-detection to relax the assumption of perfect detection on the transect line (Borchers, Zucchini and Fewster, 1998), a crucial development for many species and sampling situations (e.g. aerial surveys). Analysis of double-observer distance data is now canonically referred to as “mark-recapture distance sampling” (MRDS; Laake and Borchers, 2004) because there is a detection history (i.e. binary detection/nondetection records for each observer) in addition to recorded distances.

Several authors have investigated consequences and corrections for movement in distance sampling applications. For instance, Glennie, Buckland and Thomas (2015) showed that movement could cause considerable bias (typically positive) in distance-based abundance estimators, but did not attempt to develop methods to adjust for such bias. Hiby and Lovell (1998) developed a likelihood framework to estimate abundance when movement is random (i.e., nonresponsive to the survey platform) and occurs between successive observations.

Likewise, Borchers et al. (2010) showed that measurement errors could cause substantial (usually positive) bias in distance sampling abundance estimators. A number of authors have proposed models that account for measurement error in specific distance sampling applications (see e.g. Schweder et al., 1999; Borchers et al., 2010, and references therein).

Several observer configurations are possible within an MRDS estimation framework (Burt et al., 2014) and have important implications for bias control when animals move in response to a survey platform (i.e. “responsive” movement). In an “independent” configuration, observers detect animals independently of one another. Under this configuration it is possible to try to account for heterogeneity in detection probabilities (e.g. visual distinctiveness of different animal groups) by modelling lack of fit between the distribution of observed distances and estimated detection probabilities as a function of distance (Laake and Borchers, 2004; Borchers et al., 2006; Buckland, Laake and Borchers, 2010). The ability to account for such heterogeneity is important, since abundance estimators are negatively biased otherwise. Alternatively, in a “trial” configuration (Buckland and Turnock, 1992), one observer searches ahead, while another searches closer to the survey platform. Under this configuration, detections by the first observer are used as trials for the second observer. The trial configuration is useful for reducing bias associated with responsive movement of animals (which often positively biases abundance estimators), but one can no longer model heterogeneity in detection probability (Burt et al., 2014).

In this paper, we develop an integrated likelihood framework to account for movement and measurement error using an independent observer MRDS configuration. Specifically, we address movement between the time two observers (e.g., front and rear seat observers in aerial surveys) are able to make detections. Our objective is to account for the biasing effects of measurement error and responsive movement while also being able to model individual heterogeneity through an observer dependence specification. The remainder of this article is structured as follows. First, we describe a motivating data set, in which distance, detection histories, and individual covariates are assembled from a double-observer waterfowl aerial survey. Second, we describe a maximum marginal likelihood (MML) framework for analyzing these data. Under this framework, true animal locations are treated as latent variables. Next, we illustrate our method by analyzing the waterfowl data set and examine estimator performance with two simulation studies. We conclude with a short discussion.

## 2. Waterfowl data

In June of 2014, biologists conducted a pilot double-observer helicopter (BELL 206L on floats) survey of Arctic bird species in the Queen Maud Gulf Migratory Bird Sanctuary (Nunavut, Canada). The birds surveyed were predominantly waterfowl, but also included cranes and ptarmigan; we refer to them collectively as waterfowl for the remainder of the paper. The purpose of this particular survey was not to estimate abundance. Rather, researchers were interested in comparing estimates of detection probability from double-observer distance sampling with those from strip transects. The survey is described in greater detail elsewhere (Alisauskas and Conn, 2017), but we briefly provide information relevant to the analysis conducted later in this paper.

During the survey, two observers, one behind the other, on the same (left) side of the helicopter independently detected and recorded the perpendicular distance from the transect line to each bird group they observed. Distances were binned into 6 classes: 0-40m, 40-80m, 80-120m, 120-160m, 160-200m, and 200m+ (note that observations in the final bin are not used in subsequent analysis). They also recorded species, the number of waterfowl in each detected group (“group”), and a binary indicator for whether the waterfowl group was flapping their wings (“moving”). These data were previously analyzed by Alisauskas and Conn (2017), who used standard MRDS methods that ignored movement and measurement error in their analysis. Their analysis suggested higher detection probabilities for moving individuals, larger group sizes, and for the front seat observer (relative to a rear seat observer). They also estimated similar species effects on detection for 7 of the 9 species analyzed; here, we pool detections of these 7 species (Canada goose, king eider, long tailed duck, northern pintail, rock patarmigan, sandhill crane, and white fronted goose) to form an illustrative dataset. This protocol led to a total of 964 unique waterfowl group detections; 359 were detected by both observers, 348 by the front observer only, and 257 by the back observer only. Note that the back observer’s view of the first distance bin nearest the transect line was partially obstructed by the left helicopter float. A plot of observed distance deviations suggested asymmetrical responsive movement (away) from the aircraft for nonstationary animal groups. There were also some minor distance discrepancies for animal groups that were not moving, suggesting measurement error (Fig. 1). Hence, our objectives were to build models that formally account for movement and measurement error processes.

**FIG.1.**
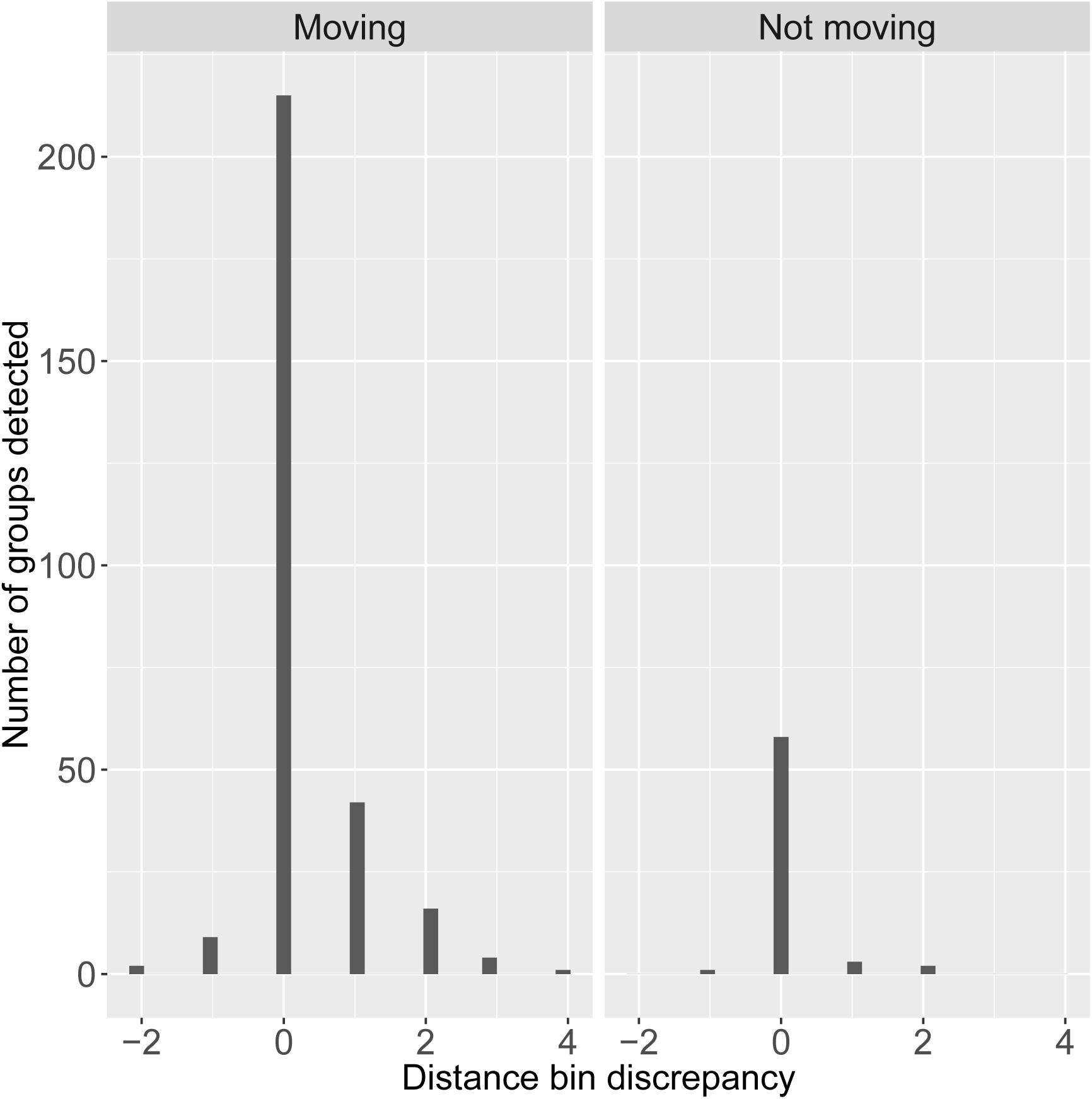
*Distribution of observed distance bin discrepancies (d_o_=2 – d_o_=1) for bird groups detected by both front (o = 1) and rear (o = 2) observers in helicopter surveys. Negative values imply movement (or measurement error) towards the helicopter, while positive values imply movement away from the helicopter. For moving birds, the distance bin observed by the rear observer tended to be further away than the bin observed by the front observer. Since the second observer always detected birds later than the front observer, this suggests that most movement was responsive away from the aircraft. For stationary birds, a nonzero distance bin discrepancy represents error in ascribing distance by either or both observers*.

## 3. Model development

Consider a double-observer MRDS survey where each observer records binned distances to detected animal groups, independently of the other observer, and a total of *n* animal groups are encountered by at least one observer (see Table 1 for a complete list of notation). We develop a two stage approach for estimating abundance in the surveyed area from such data. In the first step, a MML framework is used to simultaneously estimate parameters of detection, movement, and measurement error processes. In the second, a Horvitz-Thompson-like estimator is used to estimate abundance conditioned on parameter estimates from step 1 (a bootstrap procedure is used to quantify precision). For purposes of this paper we do not explicitly consider the problem of extrapolating abundance/density to a larger region (e.g. to unsurveyed locations), although this would be a natural extension in applied situations; we touch on this issue in the Discussion.

**TABLE 1.** *Definitions of fixed and estimated quantities for the double-observer mark-recapture distance sampling (MRDS) model incorporating movement and measurement error*

| Quantity | Definition |
| --- | --- |
| **A. Fixed quantities** |  |
| $n$ | Number of animals detected by at least one observer |
| $y_{io}$ | Binary indicator for whether animal group $i$ was detected by observer $o$ |
| $d_{io}$ | Distance bin recorded by observer $o$ for animal group $i$ (if recorded) |
| $m_i$ | A binary indicator for whether animal group $i$ was moving when observed (a single determination is made) |
| $\mathbf{x}_i$ | A vector of covariates used to explain variation in detection probability for group $i$ |
| $g_i$ | Number of animals in group $i$ (a single determination is made) |
| $\mathcal{S}$ | The set of distance bins for which data are recorded, $\mathcal{S} = \mathcal{S}_1, \mathcal{S}_2, \dots, \mathcal{S}_{n_{\mathcal{S}}}$ |
| $\mathcal{Z}$ | The set of latent distance bins used for modelling true animal locations, $\mathcal{Z} = \mathcal{Z}_1, \mathcal{Z}_2, \dots, \mathcal{Z}_{n_{\mathcal{Z}}}$ |
| $\pi_j$ | Proportion of $\mathcal{Z}$ covered by latent distance bin $j$ |
| **B. Parameters and functions of parameters** |  |
| $z_{io}$ | True (latent) distance bin of group $i$ when encountered by observer $o$ |
| $\xi_{io}$ | An indicator for whether or not observer 3 – $o$ detected group $i$ |
| $\delta_{io}$ | Perpendicular distance from the transect line to the midpoint of bin $z_{io}$ |
| $\beta$ | A vector of parameters governing logit-linear variation in detection probability |
| $\phi$ | Parameters governing animal movement |
| $\varphi$ | Parameters governing distance measurement error |
| $\theta$ | The set of detection, movement, and measurement error parameters ($\theta = \{\beta, \phi, \varphi\}$) |
| $p_{io}(z_{io})$ | Probability that observer $o$ detects group $i$ given that the group is truly in distance bin $z_{io}$ |
| $p_i^*(z_{i1}, z_{i2})$ | Probability that at least one observer detects group $i$ given the group is in distance bin $z_{i1}$ at time 1 and $z_{i2}$ at time 2 |
| $\psi(z_{i1}, z_{i2})$ | Probability that an animal that is in latent distance bin $z_{i1}$ when it passes observer 1 will be in latent distance bin $z_{i2}$ when it passes observer 2 |
| $\omega(z, d)$ | Probability that an animal group in distance bin $z$ is recorded as being in distance bin $d$ |
| $\omega(z, \mathcal{S})$ | Probability that an animal group in distance bin $z$ will have a recorded distance bin falling within $\mathcal{S}$ |
| $\mathbf{X}$ | A design matrix used to impart logit-linear structure on detection probabilities; note this will often include latent distance values, $\mathbf{z}_i$. |
| $N$ | True abundance of animals in the surveyed area |

In MRDS surveys with binned distances, observers record animals as occurring in one of 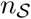 perpendicular distance bins, 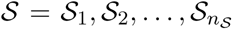. Detection probability typically decreases with distance from the transect line, and the maximum distance bin is often set such that animals further away are poorly detected and can be ignored without greatly affecting precision of abundance estimates. Movement and measurement error introduce complications: in addition to movement and measurement errors among elements of 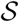, animals can potentially move into or out of 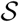, and animals outside of 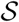 can be detected in 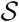. For these reasons, the models we develop rely on augmenting 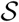 with additional distance bins to allow for movement and measurement error (Fig. 2). Call this augmented set 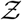.

**FIG 2.**
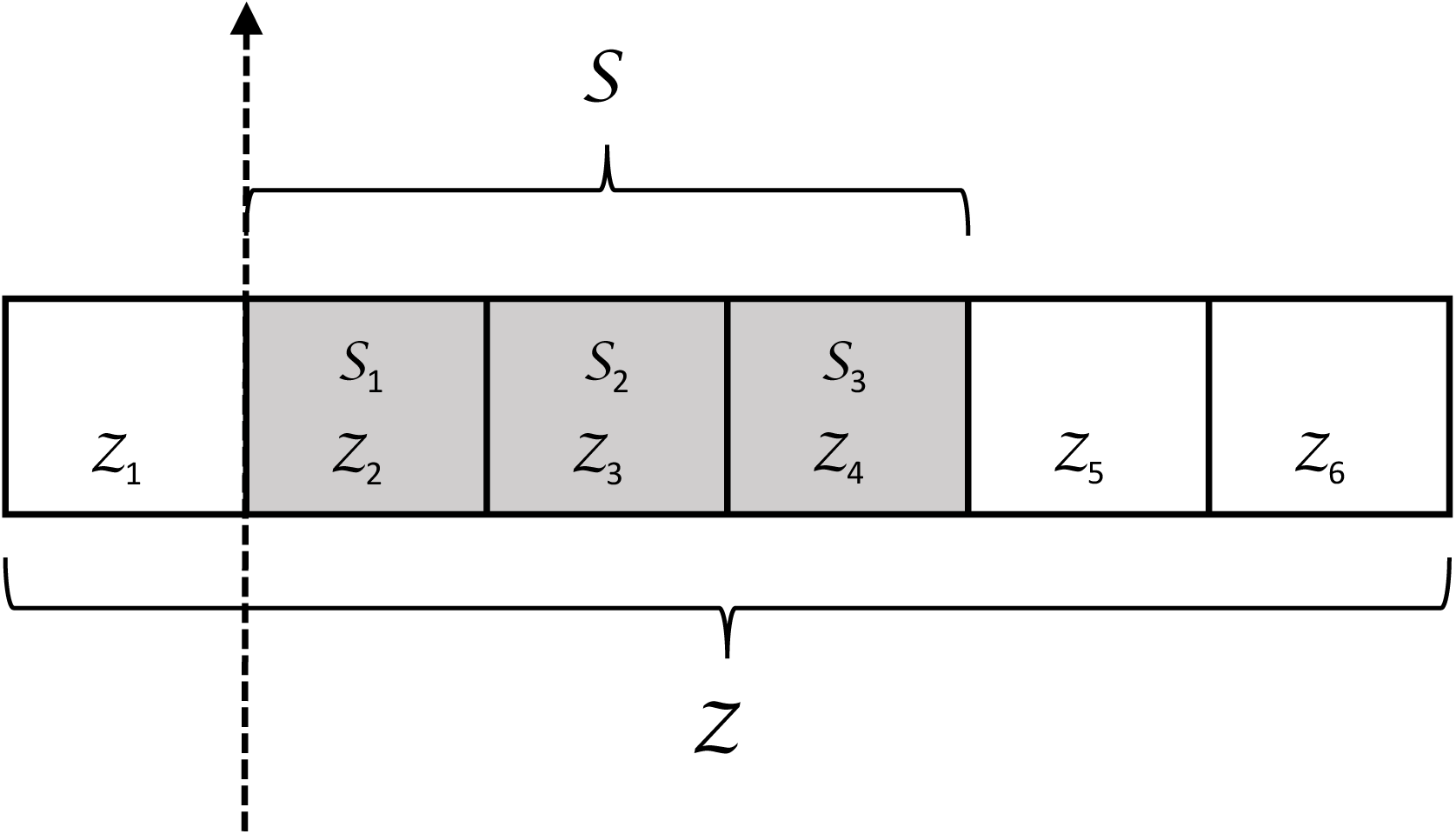
*A depiction of observed (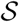) and latent (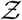) distance bins that could potentially be used in analysis of a hypothetical mark-recapture distance sampling (MRDS) survey. In this example, only animals encountered in one of the three shaded distance bins to the right of the transect line (dashed line) are recorded; however, the state space is augmented with an additional three bins to account for possible animal movement and measurement error. In practice, the number of augmented distance bins that are needed will be a function of the magnitude of the movement and measurement error processes*.

Let *y_oi_* be a binary indicator for whether or not the *i*th animal group was detected by observer *o*. Similarly, let *d_oi_* denote the distance bin recorded by observer *o* for animal group *i* (note *d_oi_* is only defined when *y_oi_* = 1). Letting bold lower case symbols denote vectors (e.g. *y_o_*. gives a sequence of detections for observer *o, i* = 1, 2,…, *n*) and bold upper case symbols denote matrices (e.g. **Y** is (2 × *n*) matrix of all detection/nondetections), we seek to define a marginal likelihood [***θ***|**Υ**, **D**, **X**], where ***θ*** = {***β***,***ϕ***,***φ***} are parameters describing probabilities of detection, movement, and measurement error, and **X** include individual covariates collected for each animal group that can be used to explain variation in detection probabilities.

## 3.1. Likelihood

To construct such a likelihood, we start with the general framework proposed by Borchers et al. (2015) for spatial mark-recapture and distance sampling surveys. Conditioning on detection, Borchers et al. (2015) suggested that the joint distribution of animal locations and detections could be written as a product of (1) a joint probability density function (pdf) for the latent locations of animals, and (2) a joint probability mass function (pmf) for the encounter histories conditional on location. We expand upon this framework to allow movement to affect the distribution of animal locations and to incorporate a measurement error mechanism.

Letting **z***_o_* denote the true locations of animals when they enter the field of view of observer *o*, we write the joint probability mass function of observed data as a product of

1. [**Ζ**|***θ***], a bivariate probability mass function for the distribution of true animal locations, given detection by at least one observer; and
2. [**Y**, **D**|**Z**, ***θ***, **X**], a model for binary detections and observed distances given true unobserved locations and individual detection covariates.

If we knew the true locations of observed animals, we could simply base inference on the likelihood

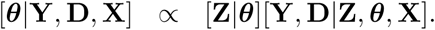

However, we do not know the actual animal locations so instead integrate (sum) over an augmented set of distance bins 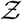 that could plausibly have resulted in a detection (see *Distribution of animal locations* for more discussion of bin augmentation). As such, we write the joint marginal likelihood of detection, movement, and measurement error parameters as

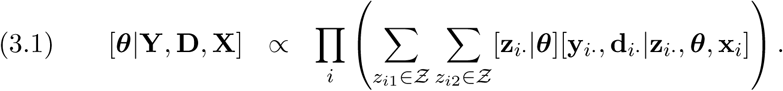

Recall that we use the *z_i_*. notation to indicate the two element vector of true distance values (over both observers’ subscripts; see Table 1 for other notation). We now describe each of the likelihood components in further detail.

## 3.1.1 Distribution of animal locations

The first component of the likelihood (Eqn. 3.1) is the joint probability mass function for the locations of group *i*, [**z***_i_*. |***θ***] given detection by at least one observer. We write this distribution as a function of (i) an initial state distribution, [*z_i_*_1_]; (ii) a movement kernel, [*z_i_*_2_|*z_i_*_1_, *ϕ*]; and (iii) detection probability by at least one observer, 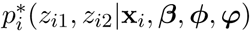. Specifically, we set

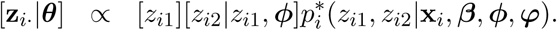

We make the assumption that the first observer (typically in a front seat) detects animal groups before movement out of the initial distance bin has occurred. Under this assumption, random placement of transect lines should help ensure that perpendicular distances of animals from the transect line are uniformly distributed in space (cf. Buckland et al., 2001). Letting *π_j_* denote the proportional diameter of distance bin *j* (i.e. 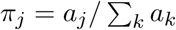 where *a_j_* is the diameter of of distance bin *j*), we simply have

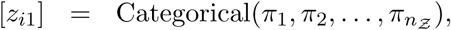

where it is understood that “Categorical” denotes a multinomial distribution with index 1, and 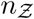 is the number of latent distance bins.

Next, the bivariate movement pmf [*z_i_*_2_|*z_i_*_1_,*ϕ*] describes the location of animal group *i* when it enters the field of view of observer 2 as a function of the location when it was in the field of view of observer 1. We model this as another categorical distribution:

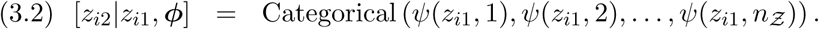

For applications in this paper, we parameterize the movement transition probabilities *ψ* using asymmetric kernels *k* (e.g. Fig 3). Using an asymmetric kernel can allow movement rates to be different toward and away from the transect line (anticipating a behavioral response to the survey platform). In particular, we set

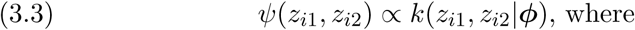

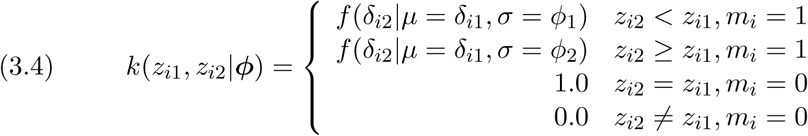

Here, *f* () gives a probability density function; in our examples, we consider Laplace (double exponential) and Gaussian distributions as choices for *f* (). Note that *δ_io_* gives the perpendicular distance from the transect line to the midpoint of distance bin *z_io_*. Also note that we assume that stationary animals (i.e. with *m_i_* = 0) do not change distance bins.

**FIG.3.**
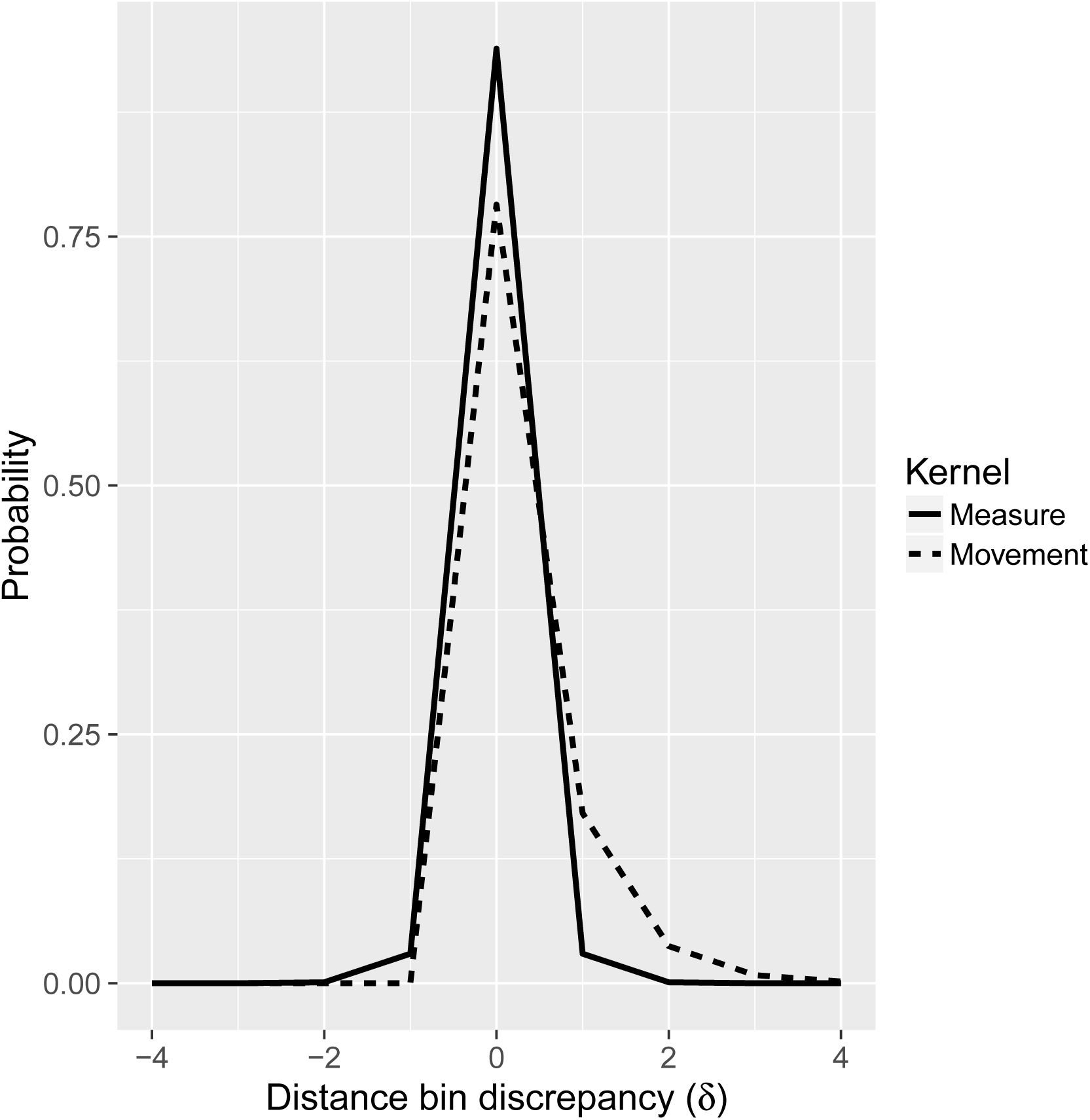
*Estimated movement and measurement error kernels for waterfowl mark-recapture distance sampling (MRDS) data from the highest ranked maximum marginal likelihood model. Measurement error used a (discretized) symmetric Laplace kernel, while movement had an asymmetric Laplace kernel*.

Finally, the thinning probability 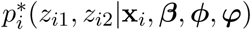 describes the probability of being detected by at least one observer for an animal that is in distance bin *z_i_*_1_ at time 1 and *z_i2_* at time 2. For generality, we calculate probability as the sum of obtaining one of the three possible detection histories: 11, 10, or 01 (detected by both observers, detected by the front observer but not the back, or detected by the back observer but not the front). In particular,

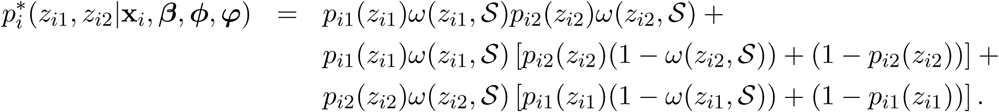

This expression is slightly different than typically encountered in mark-recapture calculus, as one must account for two ways of getting a 0 in a capture history: an observer can miss the animal group, or an observer can detect the group but determine it is out of the truncation range of the transect 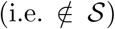. To account for the latter possibility, we make use of the measurement error kernel ***ω***, which can be parameterized similarly to *ϕ* (see Eqs. 3.3-3.4). In applications in the paper, we consider use of symmetric kernels (Gaussian or Laplace) with a single dispersion parameter, *φ*. Our expression for 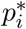 also relies on individual- and observer-dependent detection probabilities, *p_io_*(*z_io_*). In order to impart meaningful variation in detection probability, it is useful to express these in a regression framework on a logit-linear scale, such that

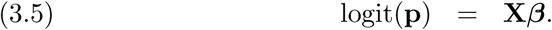

Note that we write *p_io_* as a function of *z_io_* to emphasize that the design matrix **X** will often depend on distance from the transect line (a latent quantity).

## 3.1.2 Likelihood of observed detections

The next component of the likelihood is [**y***_i_*., **d***_i_* |**z***_i_*., ***θ***, **x***_i_*], the probability of observing the particular detection history and distance bin values for animal group *i* conditional on true location. Conditional on detection by at least one observer, there are again three possible types of encounter histories: ‘11’, ‘10’, or ‘01’. For ‘11’ histories, there are 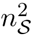 combinations of possible recorded distance bins; for ‘10’ histories, there are 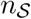 distance bins possible for observer 1; for ‘01’ histories, there are 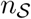 distance bins possible for observer 2. Thus, we can view [**y***_i_*., **d***_i_* |**z***_i_*., ***θ***, **x***_i_*], as a multinomial distribution with index 1 and 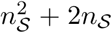 possible outcomes. The likelihood contribution for a particular animal group *i* can thus be written as

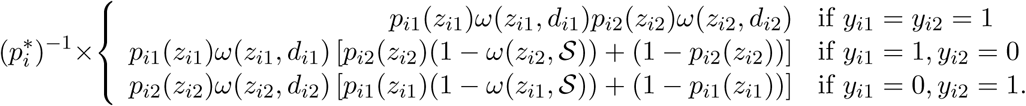

## 3.2. Horvitz-Thompson-like abundance estimator

Minimizing the negative log-likelihood in Eqn. 3.1 provides marginal maximum likelihood estimates for detection, movement, and measurement error parameters, but does not provide a direct estimate of animal abundance, *N*. We developed a Horvitz-Thompson-like procedure to calculate abundance estimates, as is common in distance sampling literature (see e.g. Buckland et al., 2004). This is especially useful when coping with detection probabilities that vary as a function of individual detection covariates, as one does not need to model covariate values for undetected animal groups. For instance, in standard MRDS applications, one might estimate abundance as

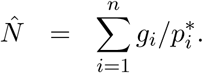

However, direct application of this estimator is clearly inappropriate under movement and measurement error, as it can potentially include animals outside of the surveyed area, or include animals that move into the surveyed area.

Since distance sampling produces estimates of abundance at a single point in time, we must first define the time and location for which the estimate applies before constructing an appropriate estimator. In the case of responsive movement away from a survey platform, we are better off referencing abundance relative to the position of animals when they enter the field of view of observer 1 than we are for observer 2 since we assume observer 1 detects an animal first. Also, since analysis only uses animals perceived to be in 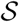, it may be best to limit the scope of inference to those animals that truly occur in 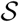. We construct a Horvitz-Thompson-like estimator for abundance in the surveyed region 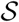 at time 1 as follows

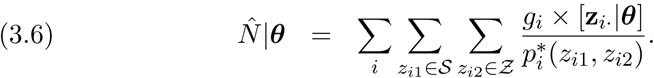

This formulation integrates over the latent position of animal groups at times 1 and 2 with the restriction that the position at time 1 is within 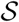.

To produce estimates of precision and confidence limits, we implemented a parametric bootstrap procedure. In particular, we approximate the sampling distribution of parameter estimates as

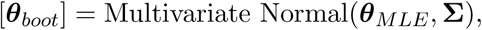

where **Σ** is a covariance matrix calculated as the inverse Hessian matrix of the likelihood evaluated at the MLE estimates. Then, for each of *k* = 1,2,…, *n_boot_* replicates, we

1. Sample ***θ****_k_* ∼ [**θ***_boot_*],
2. Transform ***θ****_k_* into real-scale parameters using inverse link functions,
3. Calculate 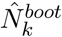 using Eqn. 3.6.

We then use quantiles of 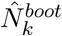 to represent confidence intervals and calculate 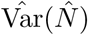 as Var(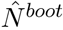).

## 3.3 Extension to incorporate detection heterogeneity

So far, we have not attempted to model detection heterogeneity outside of individual covariates (e.g. through Eqn. 3.5). However, it is common knowledge that other factors (e.g. variation in plumage, lighting, topography, background, etc.) can affect the distinctiveness of different animal groups and impart additional heterogeneity leading to (often positive) dependence in observer detection and thus negative bias in 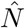 (Laake and Borchers, 2004; Buckland. Laake and Borchers, 2010; Burt et al., 2014).

In traditional MRDS applications (i.e. without movement and measurement error), one approach is to correct for this bias by estimating observer dependence parameters, typically through inclusion of an additional probability density function for observed distances in the likelihood (cf. Buckland, Laake and Borchers, 2010). However, inclusion of such a pdf in our likelihood appears problematic, as movement alters interpretation of distance distributions (Burt et al., 2014). Alternatively, MacKenzie and Clement (2016) suggested that observer dependence could also be included by modeling *conditional* detection probabilities; that is, including detection by one observer as a covariate for detection of the other. For instance, detection probabilities could potentially be written as a logit-linear function of an autocovariate ξ_iο_ = *y_i_*,_3–o_. We adapt this latter idea as a way to accommodate detection heterogeneity in data subject to movement and measurement error.

The major complication with using a detection autocovariate as a predictor in our case is that we are no longer able to say that an animal group with *y_io_* = 0 was actually undetected by observer *o*. It could, for instance, have been detected but determined to not be in 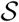. As such, we view the autocovariate ξ*_io_* as a latent variable. If *y_io_* = 1, then ξ*_i_*,_3–o_ = 1 with certainty; however, if *y_io_* = 0 we do not know whether ξ*_i_*,_3–o_ is 0 or 1. Summing over each encounter history type (11,01,or 10) subject to uncertainty about ξ*_io_*, we now need to calculate the probability of being observed by at least one observer as

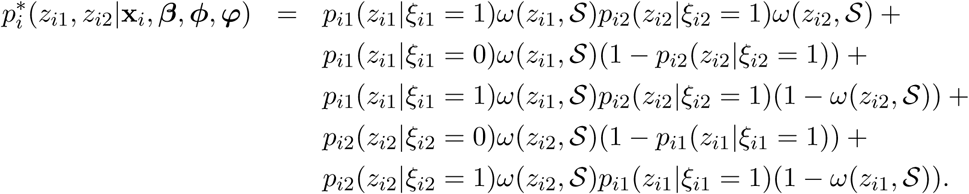

We adopt a similar construct for the observation model, [**y***_i_*., **d***_i_*|**z***_i_*·,***θ***, **x***_i_*],recasting the likelihood contribution for animal group *i* as follows according to their detection histories:

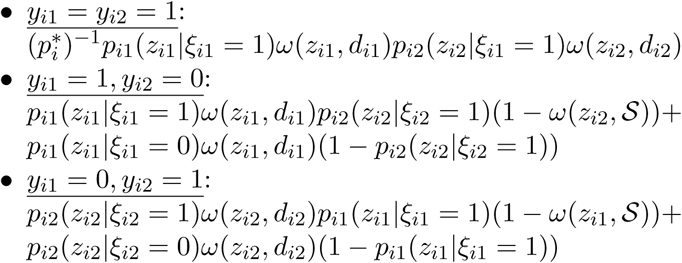

Following these adjustments, we use the “symmetric” parameterization (MacKenzie and Clement, 2016) of observer dependence to include ξ*_io_* in the logit-linear model for *p_io_*. For instance, point independence (Laake and Borchers, 2004; Buckland, Laake and Borchers, 2010), where observers are assumed to detect animal groups independently near the transect line but to have increasing dependence with distance, can be implemented by including an interaction between *z_io_* and ξ*_io_* with no main effect for ξ*_io_*. Alternatively, limiting dependence (Buckland, Laake and Borchers, 2010), where there is a base level of dependence on or near the transect line, can be implemented by including a main effect for ξ*_io_* in addition to the interaction (MacKenzie and Clement, 2016).

## 3.4. Goodness-of-fit

Goodness-of-fit is often summarized with χ^2^ tests when distance data are binned (Burnham et al., 2004). However, this depends on having adequate sample sizes and homogeneous probabilities of detection within classes of animals. This latter requirement is problematic when detection probability is written in terms of individual covariates. In order to get around this problem, we developed a simulation-based goodness-of-fit procedure similar in spirit to posterior predictive checks used in Bayesian analysis (e.g. Gelman et al., 2014). Our procedure consists of

1. Sampling ***θ****_k_* ∼ [***θ****_boot_*],
2. Simulating new data (**d***_k_*, **y***_i_*) from [**d***_k_*, **y***_k_*|**X**, ***θ***_k_].
3. Calculating a discrepancy measure *T*(**y**, **d**, ***θ***) to compare the observed data to data simulated under the model.

For instance, we might compute the proportion of observations that occur in each distance bin when subset by various explanatory variables for our observed data and compare these to the distribution of proportions that we obtain by simulating data from our model when all assumptions are met. For some specific examples, see section 4.

## 3.5. Computing

We conducted MML inference in the R programming environment (R Development Core Team, 2016). We have collated all code and data needed to recreate our analyses into an R package, `MRDSmove`. The package is currently available at https://github.com/pconn/MRDSmove/releases and will be archived on a publicly available data repository upon manuscript acceptance.

## 4. Analysis of waterfowl data

We fitted 12 MML models to our waterfowl data, varying by (1) movement and measurement kernel type (Gaussian vs. Laplace), (2) observer dependence type (none; point independence, or limiting independence), and (3) whether or not moving individuals had a different distance function than individuals that were not moving (Table 2). We calculated marginal AIC to select among these models. We also fitted two Huggins-Alho (HA; Huggins, 1989; Alho, 1990) models to our data using program MARK (White and Burnham, 1999) via an RMark (Laake, 2013) interface. The HA models suppose independent detection of observers and do not account for movement or measurement error; abundance estimates are generated with a Horvitz-Thompson-like procedure. The two HA models had the same structure but differed in how data were formatted: in the first (HA1), distance was set to *d_i_*_1_ whenever *d_i_*_1_ ≠ *d_i_*_2_; in the second (HA2), conflicting distance measurements were averaged. For the HA models, detection probability was set to the structure on the MML model with the best AIC score. All models included the following predictors within the logit-linear model for detection probability: group size, moving/not moving, observer (front vs. back), distance, distance^2^, and an interaction between the distance effects and the observer effects. The latter interaction was included because the view of the first distance bin was partially obstructed for observer 2 whose distance distribution appeared to peak further away from the helicopter (see Alisauskas and Conn, 2017).

**TABLE 2.** *Estimated abundance of waterfowl surveyed in Arctic Canada. The first 10 models account for movement and measurement error and are fitted via maximum marginal likelihood (MML), while the last two are Huggins-Alho models (HA) that ignore movement and measurement error. MML models are ranked by AIC; we also provide the number of parameters in each model (k), log likelihood (LogL) at the MMLEs, the estimated number of waterfowl groups Ĝ, and the estimated number of waterfowl (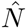). MML models varied by the functional form of movement and measurement error kernels (Gaussian vs. Laplace), the form of observer dependence (fi: full independence, pi: point independence; li: limiting independence), as well as whether the detection function included a distance: moving interaction. HA models varied by method used to reconcile distance measurements (HA1: prefer measurement of observer 1; HA2: mean distance). For HA models, only estimated bird groups are reported owing to software constraints. For reference, the number of detected bird groups was 964 and the total number of detected birds was 2666. The ‘NA’ values represent ‘not available,’ either because models did not converge, because HA likelihood and AIC values were not comparable to MML values, or because of 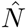 values were unavailable from the software used to conduct estimation for the HA model*.

| Model | $\Delta\text{AIC}$ | $k$ | LogL | $\hat{G}(\text{SE})$ | $\hat{N}(\text{SE})$ |
| --- | --- | --- | --- | --- | --- |
| MML.Laplace.pi.move | 0.0 | 14 | -2725.4 | 1519 (142) | 3808 (302) |
| MML.Laplace.li | 1.2 | 13 | -2727.0 | 1122 (106) | 2993 (222) |
| MML.Laplace.pi | 3.2 | 12 | -2729.0 | 1399 (99) | 3562 (209) |
| MML.Laplace.fi | 6.1 | 11 | -2731.5 | 1244 (33) | 3240 (73) |
| MML.Laplace.fi.move | 6.7 | 13 | -2729.8 | 1255 (35) | 3261 (77) |
| MML.Gaussian.pi.move | 55.5 | 14 | -2753.2 | 1516 (149) | 3800 (315) |
| MML.Gaussian.li | 56.5 | 13 | -2754.7 | 1130 (118) | 3009 (247) |
| MML.Gaussian.pi | 58.3 | 12 | -2756.6 | 1394 (101) | 3552 (212) |
| MML.Gaussian.fi | 62.9 | 11 | -2760.0 | 1248 (35) | 3248 (80) |
| MML.Gaussian.fi.move | 63.6 | 13 | -2758.2 | 1259 (35) | 3269 (79) |
| MML.Gaussian.li.move<sup>†</sup> | NA | NA | NA | NA | NA |
| MML.Laplace.li.move<sup>†</sup> | NA | NA | NA | NA | NA |
| HA1 | NA | 10 | NA | 1239 (47) | NA |
| HA2 | NA | 10 | NA | 1278 (56) | NA |

<sup>†</sup> Did not converge

AIC strongly favored models with Laplace movement and measurement error kernels (Fig. 3) over Gaussian kernels, although the impact of the functional form of the kernel on resultant abundance estimates was quite small (Table 2). The highest ranked model had an interaction between distance and moving/not moving, suggesting different detection function shapes for moving vs. stationary animals. However, pairwise model comparisons with and without such an effect had similar AIC scores, so this effect was likely small (also see Fig. 4). Point independence (‘pi’) and limiting independence (‘li’) models were favored over full independence (‘fi’) models, suggesting some level of detection heterogeneity that was not captured via gathered covariates.

**FIG.4.**
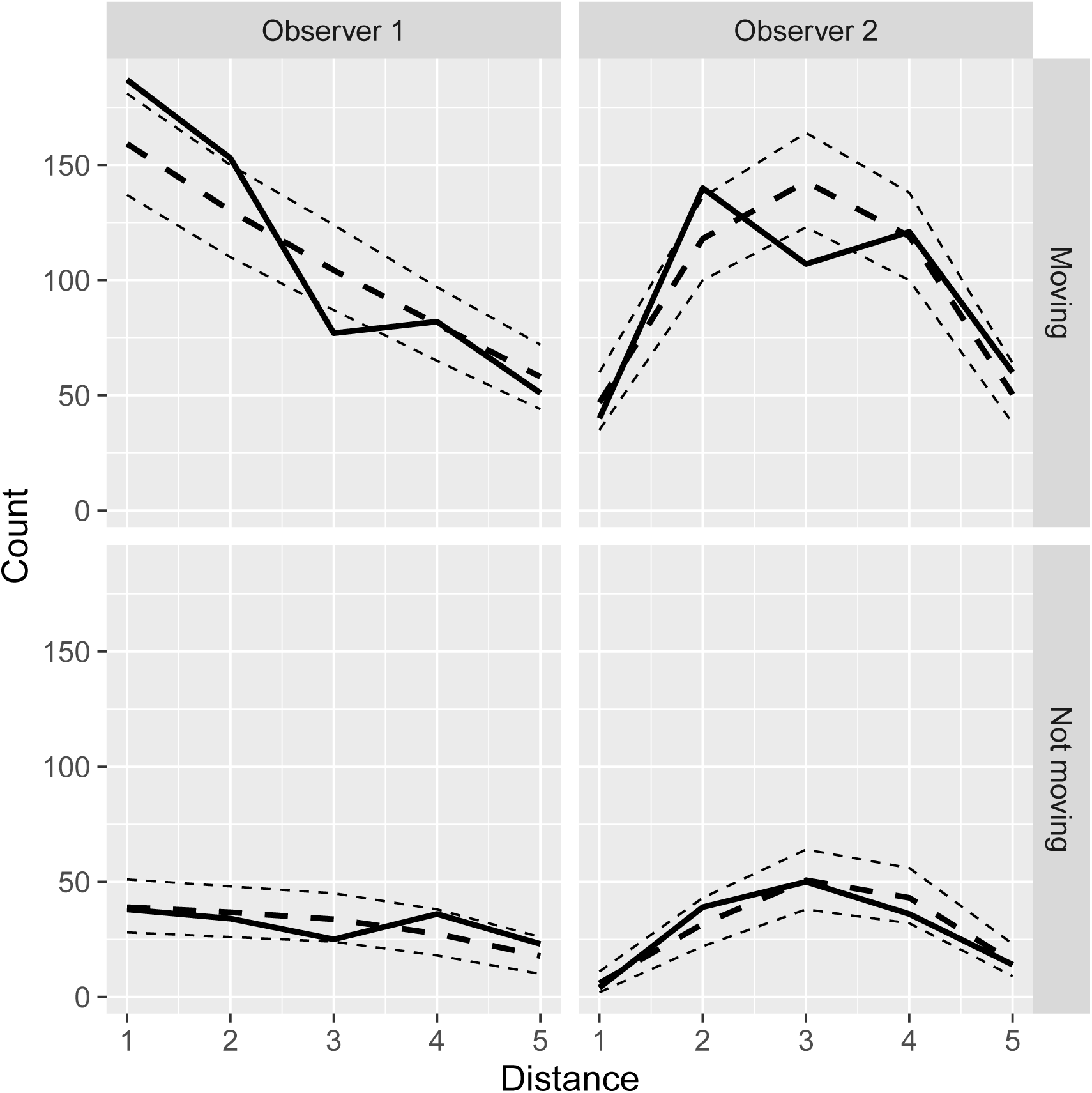
*A plot of the number of observed and predicted waterfowl groups by observer and movement status. Observed data are given by the thick solid line, while the thick dashed line represents mean predictions and the thin, dashed lines represent 2.5th and 97.5th quantiles of model-based simulations (including variance due to uncertainty of MML estimates*).

The form of dependence had large effects on abundance estimates and accompanying standard errors. In general, ‘li’ models produced the smallest estimates (*Ĝ*= 1122 to 1130), ‘fi’ models produced the next highest estimates (*Ĝ* = 1244 to 1259), and ‘pi’ models produced the highest estimates (*Ĝ* = 1394 to 1519; Table 2). Models with similar support often produced estimates of abundance that were quite different. For instance, the top two models (including a pi model and an li model) were only 1.2 AIC units apart but produced estimates of *Ĝ*= 1519 and *Ĝ*= 1122, respectively. The ‘pi’ models predict increasing observer dependence with distance, while the ‘li’ models suggested strongly negative observer dependence near the transect line which linearly increased to positive dependence in distance bin 5 and beyond. This latter type of observer dependence could occur if observers have different fields of view and are likely to detect different animal groups close to the aircraft, but are more likely to see the same animals (presumably the highly distinctive ones) farther away.

Plots of movement and measurement error kernels (Fig. 3) for the highest ranked model resembled raw data histograms (Fig. 1). However, inclusion of movement and measurement error in the model did not appear to largely affect abundance estimates. For instance, HA1 and HA2 (the models without movement or measurement error) produced estimates of 1239 and 1278 waterfowl groups, respectively. By comparison, the 4 ‘fi’ models (which, like the HA models, presume conditional independence in observer detections), produced estimates of 1244-1259. In our example, it seemed far more important to account for different types of observer dependence. If an estimate were needed for management or conservation purposes, it would be wise to compute a model averaged estimate that incorporates uncertainty about the correction functional form of observer dependence (as well as attendant standard errors, which are much higher for ‘li’ and ‘pi’ models than for ‘fi’ models). We note that several of the ‘li’ models did not converge, a relatively frequent occurence when fitting MRDS models (Buckland, Laake and Borchers, 2010; MacKenzie and Clement, 2016).

To examine fit of our model to the data, we compared the properties of our MRDS dataset to 1000 data sets simulated from the highest ranked AIC model. In general, data sets simulated under our model had similar proportions of animals observed in the five distance bin classes as we observed in the field (Fig. 4). A notable exception was a tendency to overpredict the proportion of moving animals in distance bin 3. We are unsure why there may have been a dip in detections in the third distance bin, but have resisted the urge to consider more highly parameterized structures since a smooth decrease in the number of animals encountered as a function of distance is often expected a priori (Buckland et al., 2001), and it would be difficult to fit this particular “dip” in our distance data without making the detection model multimodal. Our model did a reasonable job in replicating the proportions of animals with each detection history type observed in the field. For instance, the number of ‘11’, ‘10’ and ‘01’ histories compiled for moving animals was 289, 261, and 179, respectively; these compared to 95% simulation intervals of (257,307), (227,276), and (173,219). For stationary animals, we observed 64 ‘11’, 92 ‘10’ and 79 ‘01’ histories compared to simulation intervals of (53,80), (74,103), and (68,95).

## 5. Simulation studies

We conducted two simulation studies to investigate bias, precision, and confidence interval coverage of our MML models and compared these to other MRDS analyses that do not account for movement and measurement error. The first simulation study assumed independence between observer detections (i.e., no residual detection heterogeneity). The second experiment focused on performance of different approaches to estimation when heterogeneous detection probabilities were simulated using random effects.

## 5.1 Simulation study I: Basic model performance

Our first simulation study was designed to investigate estimator performance over different movement and measurement error rates, and only considering variation imparted by measurable covariates. For this study, we simulated three different Gaussian movement kernel (Eqn. 3.4) scenarios, corresponding to (i) no movement (*ϕ*_1_ = *ϕ*_2_ = 0), (ii) symmetric movement (*ϕ*_1_ = *ϕ*_2_ = 0.7), and (iii) asymmetric movement with much higher rates of movement away from the transect line than towards the transect line (*ϕ*_1_ = 0.5, *ϕ*_2_ = 1.5). We considered two levels of measurement error for each movement scenario: no measurement error, or minor measurement error (*φ* = 0.5). The latter value of measurement error was chosen to approximate the level of error we observed in our waterfowl data.

In each of 500 simulations for the 6 movement and measurement error scenarios, we conducted the following steps:

1. For each of *i* ∈ 1, 2,…, 1000 animals, we simulated an initial, latent position *Z*_*i*1_ in 10 equally sized distance bins using a uniform distribution.
2. After generating *m_i_* ∼ Bernoulli(0.75) (so that approximately 75% of animals were moving), we simulated *Z*_*i*2_ using Eqn. 3.2. For animals with *m_i_* = 0, we simply set *Z*_*i*2_ = *Z*_*i*1_.
3. We simulated *y_io_* and *d_io_* using detection and measurement error models, where the first five distance bins were subject to observation (i.e. 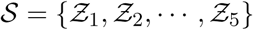). Detection probabilities were configured as

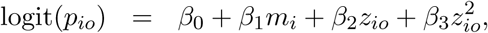

where *β*_0_ = 1, *β*_1_ = 0.5, *β*_2_ = 0.07, and *β*_3_ = −0.09.
4. We fit a sequence of three models to each such data set. These included (i) the movement and measurement error model proposed in this paper (configured with 8 latent distance bins), as well as the two Huggins-Alho models described in section 4. For all three estimation procedures, we used the same structure when estimating *p_io_* as used to generate the data. For simulations where data were generated with *ϕ* = 0 or *φ* = 0, we fixed the corresponding parameter in the estimation model to zero to prevent numerical errors.
5. For each model and data set, we tabulated bias in abundance (note that 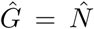 since group sizes were all 1.0), coefficient of variation (CV), 95% confidence interval coverage, and root mean square error (RMSE).

Note that in initial simulation work, we also fit movement and measurement error models with 10 latent distance bins, finding that results were almost identical to those with 8 latent distance bins (parameter estimates were often within 0.0001 of each other).

In general, bias from our new method was zero to slightly negative, while positive bias from the HA models could be substantial when movement and/or measurement error occurred (up to 10%; Table 3). Precision and mean squared error were always better for the MML models than the HA models, with confidence interval coverage closer to nominal for most of the MML to HA model comparisons. However, coverage was less than nominal (85-91% for a 95% interval) for the MML models, suggesting that our bootstrap-based interval estimation procedure produced estimates of variance that were too small. Interestingly, HA1 estimates tended to have better properties (lower bias, better coverage, lower RMSE) than HA2 estimates, suggesting that taking distance values from observer 1 may be a better strategy than averaging distance values to resolve discrepancies if one cannot model movement and measurement error directly.

**TABLE 3.** *Mean proportion relative bias (RelBias), coefficient of variation (CV), 95% confidence interval coverage (Cover), and root mean squared error (RMSE) for the two simulation studies. For the first simulation scenario, “Configuration” gives values for movement (σ_1_ and σ_2_) and measurement error (φ) parameters, e.g. (0,0,0), respectively; in simulation study 2, it indicates these parameters as well as expected population size in the surveyed area N = 200 or N = 1000. Three estimation models (Model) were fitted to each data set in simulation study 1: the maximum marginal likelihood (MML) model accounting for movement and measurement error, and two Huggins-Alho models which do not account for movement, measurement error, or observer dependence (HA1 and HA2; described in the text). For simulation scenario two, we fitted an additional MML model that accounts for observer dependence (MMLd)*.

| Configuration | Model | RelBias | CV | Cover | RMSE |
| --- | --- | --- | --- | --- | --- |
| **A. Simulation study 1** |  |  |  |  |  |
| (0,0,0) | MML | 0.00 | 0.03 | 0.91 | 214 |
| (0,0,0) | HA1 | 0.01 | 0.04 | 0.95 | 532 |
| (0,0,0) | HA2 | 0.01 | 0.04 | 0.95 | 529 |
| (0.7,0.7,0) | MML | 0.01 | 0.03 | 0.87 | 268 |
| (0.7,0.7,0) | HA1 | 0.04 | 0.05 | 0.84 | 1144 |
| (0.7,0.7,0) | HA2 | 0.06 | 0.05 | 0.80 | 1763 |
| (0.5,1.5,0) | MML | -0.02 | 0.03 | 0.89 | 371 |
| (0.5,1.5,0) | HA1 | 0.08 | 0.07 | 0.77 | 3086 |
| (0.5,1.5,0) | HA2 | 0.07 | 0.07 | 0.84 | 2883 |
| (0,0,0.5) | MML | 0.00 | 0.03 | 0.91 | 245 |
| (0,0,0.5) | HA1 | 0.01 | 0.05 | 0.95 | 490 |
| (0,0,0.5) | HA2 | 0.03 | 0.05 | 0.93 | 740 |
| (0.7,0.7,0.5) | MML | 0.01 | 0.03 | 0.87 | 306 |
| (0.7,0.7,0.5) | HA1 | 0.04 | 0.05 | 0.86 | 1237 |
| (0.7,0.7,0.5) | HA2 | 0.07 | 0.06 | 0.75 | 2676 |
| (0.5,1.5,0.5) | MML | -0.03 | 0.03 | 0.85 | 471 |
| (0.5,1.5,0.5) | HA1 | 0.07 | 0.07 | 0.82 | 3045 |
| (0.5,1.5,0.5) | HA2 | 0.10 | 0.08 | 0.77 | 4387 |
| **B. Simulation study 2** |  |  |  |  |  |
| (0,0,0), $N = 200$ | MMLd | 0.03 | 0.12 | 0.94 | 435 |
| (0,0,0), $N = 200$ | MML | -0.04 | 0.05 | 0.89 | 194 |
| (0,0,0), $N = 200$ | HA1 | -0.06 | 0.06 | 0.83 | 312 |
| (0,0,0), $N = 200$ | HA2 | -0.06 | 0.06 | 0.83 | 312 |
| (0,1.5,0.5), $N = 200$ | MMLd | -0.01 | 0.15 | 0.96 | 692 |
| (0,1.5,0.5), $N = 200$ | MML | -0.08 | 0.06 | 0.74 | 440 |
| (0,1.5,0.5), $N = 200$ | HA1 | 0.04 | 0.13 | 0.95 | 2581 |
| (0,1.5,0.5), $N = 200$ | HA2 | 0.00 | 0.12 | 0.92 | 921 |
| (0,0,0), $N = 1000$ | MMLd | 0.03 | 0.05 | 0.91 | 2574 |
| (0,0,0), $N = 1000$ | MML | -0.05 | 0.02 | 0.33 | 3519 |
| (0,0,0), $N = 1000$ | HA1 | -0.08 | 0.02 | 0.22 | 6315 |
| (0,0,0), $N = 1000$ | HA2 | -0.08 | 0.02 | 0.22 | 6312 |
| (0,1.5,0.5), $N = 1000$ | MMLd | -0.03 | 0.05 | 0.94 | 3092 |
| (0,1.5,0.5), $N = 1000$ | MML | -0.09 | 0.02 | 0.06 | 8817 |
| (0,1.5,0.5), $N = 1000$ | HA1 | -0.01 | 0.05 | 0.92 | 2607 |
| (0,1.5,0.5), $N = 1000$ | HA2 | -0.03 | 0.04 | 0.84 | 3080 |

## 5.2 Simulation study II: Heterogeneous detection

In our second simulation scenario, we examined performance of our proposed approach when MRDS data are simulated with highly heterogeneous detection probabilities. The main structure of our simulations was largely similar to the preceding section. We considered two different movement and measurement error scenarios corresponding to none (*ϕ*_1_ = *ϕ*_2_ = *φ* = 0) and to movement away from the survey line (*ϕ*_1_ =0, *ϕ*_2_ = 1.5, *φ* = 0.5). For each of these scenarios, we considered two different expected sample sizes in the sampled area: *E*(*N*) = 200 and *E*(*N*) = 1000. In each combination of simulation replicates, we conducted 500 simulations via following steps:

1. For each of *i* ∈ 1,2,…, 2*E*(*N*) animals, we simulated an initial, latent position *z_i_*_1_ in 10 equally sized distance bins using a uniform distribution.
2. We generated *m_i_* and *z_i_*_2_ as in Simulation Study 1.
3. We simulated *d_io_* and *y_io_* as in simulation study 1, once again using 5 observable distance bins. However, we used a half-normal model for detection probability,

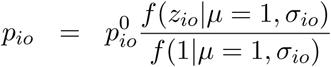

where 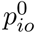 gives detection probability in the first distance bin, and the half normal model describes how detection probability declines in bins that are farther away. These models were further parameterized as

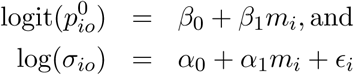

where *β_ο_* = *α*_0_ = 1, *β*_1_ = 0.5, *α*_1_ = 0.2, and *∊_i_* ∼ Uniform(–0.7,0.7). The half-normal model seemed a reasonable way to implement point independence (Laake and Borchers, 2004) using random effects (Fig.5).
4. We fitted four models to each such data set. These included the same three models from Simulation Study 1, and a fourth, marginal likelihood model that attempted to estimate an observer dependence parameter in addition to movement and measurement error. Observer dependence used a point independence specification (i.e. an interaction between *ξ_io_* and *δ_io_*).
5. For each model and data set, we tabulated bias, coefficient of variation (CV), 95% confidence interval coverage, and root mean square error (RMSE).

Simulations suggested that the MML model with observer dependence did a reasonable job at estimating abundance under all scenarios (Table 3) even though the estimation model differed from the data generating model (polynomial vs. half normal detection model; observer dependence effect vs. random effects). In particular, bias was low (-0.03 to 0.03) and 95% confidence interval coverage was close to nominal (0.91 - 0.96) for all scenarios examined. In contrast, bias of models ignoring observer dependence could be considerable (up to -9%) with precision that was too high, leading to confidence interval coverage that was too low (as low as 6% in one scenario). Not surprisingly, bias was typically negative when ignoring observer dependence. However, there was a mediating effect on bias whenever data were simulated subject to both movement, measurement error, and observer dependence. Since movement and measurement error alone induce positive bias, and observer dependence alone produces negative bias, both processes combined attenuated bias. For instance, HA models actually performed better when both sources of bias were present than when one source of bias was present.

**FIG.5.**
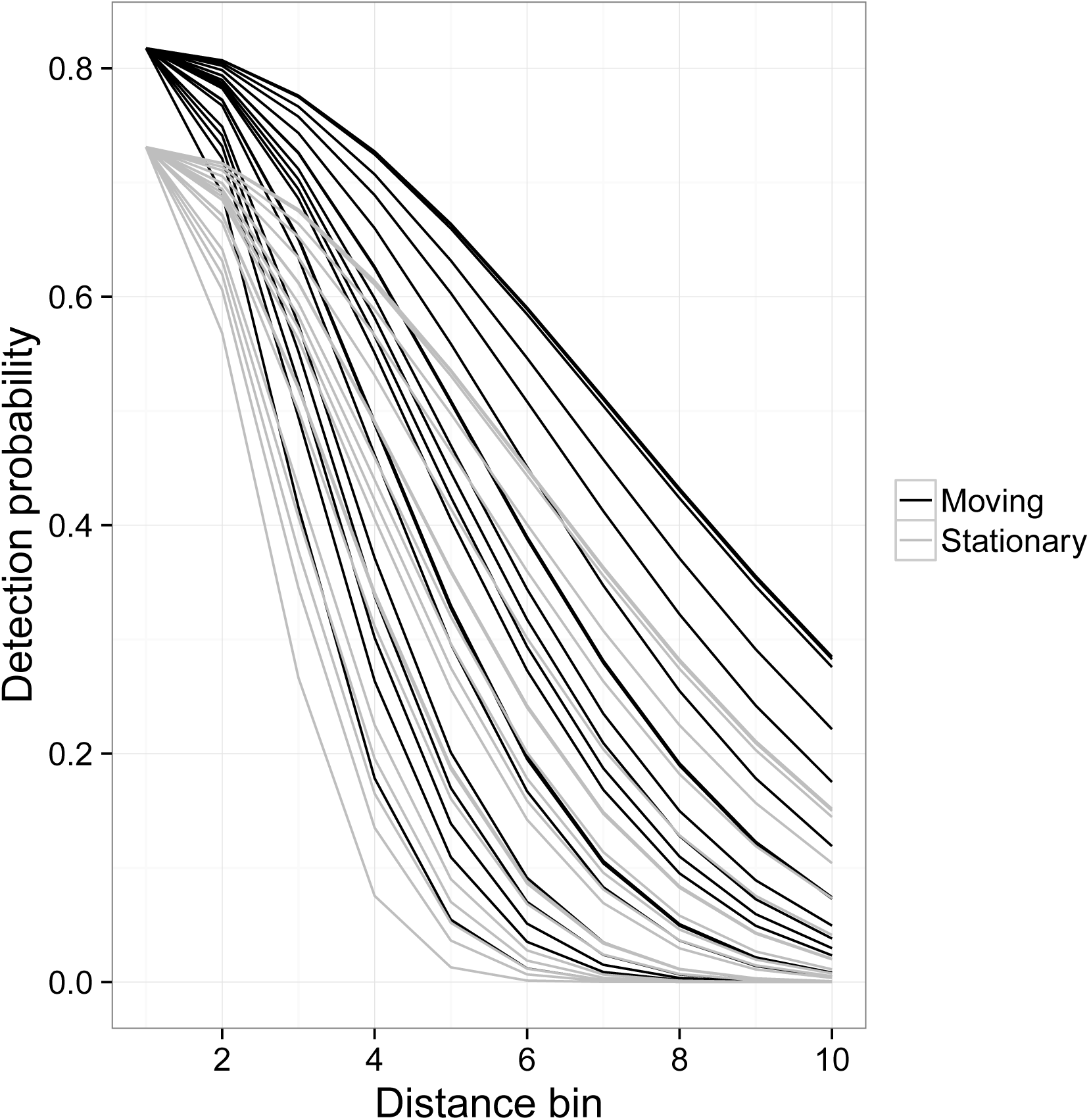
*Detection probability for a random sample of 20 individuals from Simulation Study 2, where heterogeneity is incorporated via a random effect on the log of the standard deviation associated with a half-normal detection model. Detection probabilities are presented for cases where animals are moving (black lines) or not moving (gray lines)*

## 6. Discussion

In this paper, we developed an approach to account for movement and measurement error in MRDS analyses when observers independently record distances to animals, and when there is a binary covariate for movement. In simulation studies, our approach exhibited low bias and RMSE when compared to a procedure that ignores movement and measurement error (the latter resulted in positive biases of up to 10%). Importantly, we were able to conduct estimation even in the face of residual detection heterogeneity, which seems like a useful advance. Indeed, estimation of abundance in our field study was much more sensitive to different functional forms for observer dependence than it was to different functional forms for movement or measurement error.

Several avenues of future research are desirable. First, our bootstrap-based estimates of variance resulted in confidence interval coverage that was less than nominal in some of the simulation scenarios. A more robust method for producing confidence intervals for Horvitz-Thompson-like abundance estimates would be useful. Second, although our focus here was on errors in distances, other errors may occur (e.g. errors in group size determinations, individual covariates, species, etc.). Errors in species identification can be particularly problematic (see e.g. Conn et al., 2014) and should ultimately be addressed in multi-species surveys. Third, we have assumed additive measurement error in the present development; in some situations, multiplicative measurement error (whereby animals further away are subject to greater measurement error; Borchers et al., 2010) may make more sense. Finally, we have temporarily ignored the problem of expanding estimates from the surveyed area to some larger area of inference. One approach to expanding the scope of inference would be to include a sample inclusion probability in the denominator of the Horvitz-Thompson estimator (i.e. Eqn. 3.6). Another approach would be to produce sequences of estimates for different surveyed areas (presumably sharing detection and movement/measurement error parameters between areas), and to use such estimates as responses for subsequent spatial modelling efforts (see e.g. Miller et al., 2013).

In this paper we conditioned on binary variables *m_i_* for whether a detected group was moving or not. This approach let us separately estimate movement from measurement error by making the assumption that animals with *m_i_* = 0 do not move. In other situations and study taxa (e.g. many marine mammals), all animals may be moving in some fashion, and thus there may be insufficient data to separate these processes. In these circumstances, auxiliary data (e.g. animals with known location to estimate measurement error; cf. Borchers et al., 2010) may be needed to implement our methods.

One exciting avenue for future research would be to expand our type of modelling approach to allow movement within spatial capture-recapture (SCR; see e.g. Borchers and Efford, 2008; Royle et al., 2013) models. The generalized likelihood structure of MRDS and SCR is actually very similar (Borchers et al., 2015; Borchers and Marques, 2017), so incorporating movement could likely be accomplished using the same construct in the paper (i.e. by viewing an animals’ locations as unobserved latent variables and integrating over all possible sequences of locations). The challenge would likely be a numerical one, as space would need to be increased from one to two dimensions and over a finer mesh, and the temporal dimension would need to increase from two observers to a finite number of sampling occasions. One approach to high dimensional integration would be to adopt a Bayesian perspective within a data augmentation framework (Royle, Dorazio and Link, 2007; Conn, Laake and Johnson, 2012).

## Acknowledgements

We thank Jeff Laake and Brett McClintock for comments on an earlier draft of this paper.

## DISTANCE SAMPLING WITH MOVEMENT

7600 SAND POINT WAY NE SEATTLE, WA 98115 USA E-MAIL:
PRAIRIE AND NORTHERN WILDLIFE RESEARCH CENTRE 115 PERIMETER RD. SASKATOON, SK S7N 0X4 CANADA E-MAIL

